# Translational inhibition and phase separation primes the epigenetic silencing of transposons

**DOI:** 10.1101/2020.04.08.032953

**Authors:** Eun Yu Kim, Ling Wang, Zhen Lei, Hui Li, Wenwen Fan, Jungnam Cho

## Abstract

Transposons are mobile DNAs that can cause fatal mutations. To counteract these genome invaders, the host genomes deploy small interfering (si) RNAs to initiate and establish the epigenetic silencing. However, the regulatory mechanisms for the selective recognition of transposons by the host genomes remain still elusive. Here we show that plant transposon RNAs undergo frequent ribosome stalling caused by their inherently unfavourable codon sequence usage. The ribosome stalling then causes the RNA truncation and the localization to siRNA bodies, which are both critical prerequisites for the siRNA processing. In addition, SGS3, the key protein in the siRNA biogenesis pathway, forms liquid droplets *in vitro* through its prion-like domains implicating the role of liquid-liquid phase separation in the formation of the siRNA bodies. Our study provides a novel insight into the regulatory mechanisms for the recognition of invasive genetic elements which is essential for the maintenance of genome integrity.

## Introduction

Transposable elements (TEs, transposons) are mobile genetic elements that jump around the genomes, and thus pose a significant threat to genome integrity^1,2^. The host genomes have evolved elaborate mechanisms involving various kinds of short interfering (si)-RNAs to battle against these genomic parasites^2–4^. In plants, the 24-nucleotide (nt) siRNAs are the predominant form of siRNAs and mediate the RNA-directed DNA methylation (RdDM) that maintains the epigenetic silencing and heterochromatin formation of transposons^5^. While the canonical RdDM is mainly for reinforcing the silent state of transposons, the recently proposed alternative RdDM pathway suppresses the active transposons that are induced upon the epigenetic mutations and in several development stages^3,6–10^. RNA-DEPENDENT RNA POLYMERASE 6 (RDR6)-SUPPRESSOR OF GENE SILENCING 3 (SGS3) complex serves as a detector of transposon RNAs that selectively processes them into 21 or 22-nt siRNAs (also referred to as epigenetically activated siRNAs, easiRNAs)^11,12^. Transposon-derived siRNAs trigger both post-transcriptional repression by cleaving the transposon RNAs (and then subsequently initiating the secondary siRNA biogenesis) and establishment of the epigenetic silencing by recruiting DNA methyltransferases to the target TE chromatin^5,7,8^. The biogenesis of 21 or 22-nt siRNAs in plants is initiated by RDR6 which is an RNA-dependent RNA polymerase that forms doublestranded RNAs by templating directly the target RNAs^13^. SGS3 is an RNA-binding protein that interacts with RDR6 and is essential for its function^13^. The duplex RNAs are then sliced to 21 or 22-nt siRNAs by DICER-LIKE 3 or 4 (DCL3/4)^8,14,15^. Since RDR6 templates are directly derived from the RNAs of those mobile elements, RDR6 and the resulting easiRNAs are deemed the first line of the host immune system against the invasive genetic elements.

The RDR6-mediated easiRNA production pathway (RDR6-RdDM) is usually prevented in the transcripts derived from genes by the RNA decay pathways^16–18^. This raises an important question of how RDR6 specifically recognizes its TE targets and establish their epigenetic silencing. Previous reports showed that the initial cleavage of target transcripts is a critical prerequisite for RDR6 recognition^19,20^, however, the precise mechanisms for the initial RNA cleavage prior to easiRNA production is still unclear. Several studies attempted to answer this question by suggesting that miRNA-mediated^4,8^ or Nonsense-Mediated RNA Decay (NMD) pathway^21^ induces the initial cleavage of a subset of transposon RNAs. Nonetheless, the exact cellular phenomena happening to RNAs destined to the easiRNA biogenesis pathway is yet to be known.

The specificity of the easiRNA production pathway is also provided by spatial confinement into siRNA bodies where RDR6 and SGS3 are localized. Unfortunately, how the target TE RNAs are funnelled to the siRNA bodies are unknown. Interestingly, the siRNA bodies often colocalize with the stress granules, the membrane-less cytoplasmic organelles that are formed by OLIGOURIDYLATE-BINDING PROTEIN 1 b (UBP1b)^22^. UBP1b is one of the plant homologs of T-cell Intracytoplasmic Antigen 1 (TIA-1), a viral translational repressor in mammals^23^, and is known to suppress the translation of transposons in *Arabidopsis^24^.* A recent report has shown that stress granules are rich with weakly translating mRNAs in yeast and human^25^, further supporting the notion that stress granules are associated with the translational repression. This hints at the potential linkage of weak translation and siRNA body localization of transposons which can provide additional selectivity to TE RNAs for easiRNA production.

Cellular compartmentalization is a common biological phenomenon that enhances the efficiency and specificity of certain cellular pathways. Liquid-liquid phase separation (LLPS) of ribonucleoproteins has recently emerged as a potent physiochemical driver for such compartmentalization and is relevant to diverse cellular processes and human diseases^26,27^. LLPS usually happens to proteins containing the prion-like domains or low complexity domains. Stress granule is one of the best-known cytoplasmic bodies that is formed through LLPS in human cells^28–30^. Despite the strong conservation of the prion-like domains, however, the plant components of stress granules and their accompanying siRNA bodies have never been assessed so far for their abilities to phase separate *in vitro.*

Previously, we reported that CpG dinucleotide-rich sequences exhibit epiallelic behaviour, i.e. they are less capable of regaining DNA methylation once lost^31^. In contrast, CpG-scarce sequences that are usually found in transposons readily regain DNA methylation presumably through the siRNAs originated from TE RNAs^31^. We then further extended our investigation to better understand the cellular events that connect the sequence bias of TEs to the siRNA biogenesis. In this study, we suggest that transposons are different from genes in the codon sequence usage and is rich with the codons that are unfavourable for translation. The weak translation then serves as a signal that leads to RNA truncation and siRNA body localization of transposons which are essential for easiRNA biogenesis. Besides, we demonstrate that the formation of siRNA bodies is mediated by LLPS of SGS3, implicating that phase separation is important for RDR6-RdDM pathway. Our work uncovers the general features of mobile genetic elements that are selectively guided to the siRNA biogenesis pathway, which is critical for maintaining the genome integrity.

## Results

### Codon sequence bias and reduced translation efficiency of transposons

Similar to the sequence features of high CpG density associated with the epiallelic loci in *Arabidopsis^3^,* several other studies also showed that GC3 contents (GC contents at the third nucleotide positions of codons) are negatively correlated with siRNAs and DNA methylation levels^32–34^. Besides, it has been also suggested that the expression of transposons by RNA Polymerase II is essential for the siRNA production and *de novo* DNA methylation^7,12^. These indicate that the unique epigenetic behaviour of transposons might be attributed to their RNA sequence feature, however, its mechanistic understanding is still largely lacking. In order to dissect the mechanisms underlying the selective recognition of TEs by the host genomes for the initiation of epigenetic silencing, we first thoroughly interrogated the base composition of coding sequences in the rice genome. As shown in Fig. 1a, while the GC contents at the first and second nucleotide positions of the codons (indicated as GC1 and GC2, respectively) are similar between genes and transposons, the GC3 contents of transposons are remarkably lower than those of genes. Such divergence of codon sequence usage might contribute to weak translation of TE transcripts that was partly suggested previously for rice^35^. We paid attention particularly to translational repression of TEs because it often induces RNA cleavage through the so called No-Go RNA Decay (NGD) pathway^36–40^. In addition, the core NGD complex Pelo-Hbs1 was previously reported to suppress transposon activity in *Drosophila^41^.* Given that RNA truncation is an essential prerequisite for RDR6 targeting and subsequent easiRNA biogenesis, we hypothesized that the TE RNAs might be more prone to cleavage caused by the reduced translation. In order to obtain the transcriptome-wide view of transposon translation, we analysed the public translatome data (PRJNA298638) generated from rice callus^35^. We chose rice callus because *in vitro* tissue cultured callus samples contain more active transposons^42^. By assessing the translational efficiency index (TEI) defined as the relative level of translation to transcription, we observed that rice TEs are significantly weaker in translation than genes (Fig. 1b and c).

**Figure 1.**
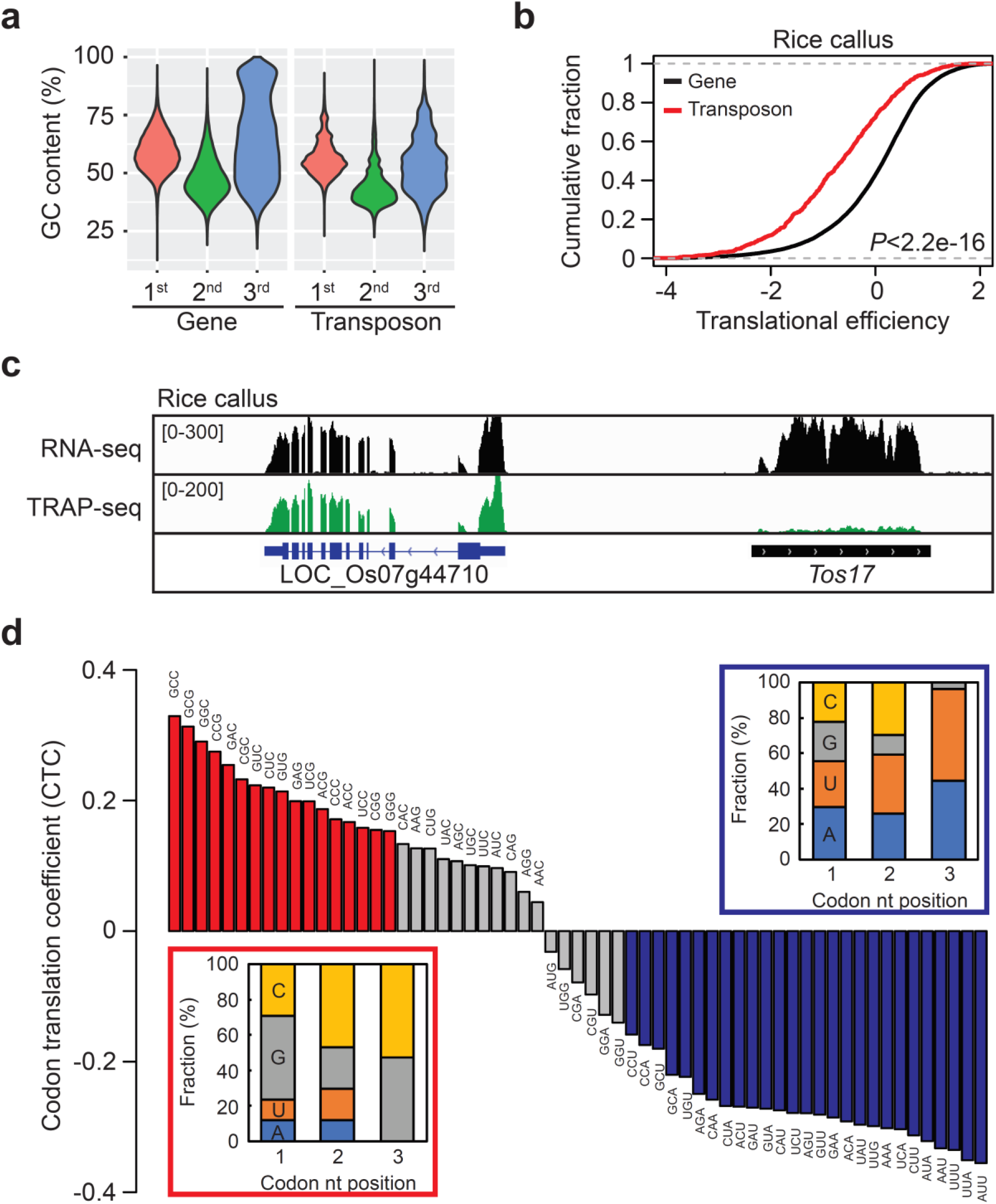
Reduced translational efficiencies of transposons in rice. **a**, GC contents of genes and transposons of rice shown for different codon nucleotide positions. **b**, Translation efficiency of genes and transposons. Translational efficiency indices (TEI) are determined as the log2-ratio of TRAP-seq (Translating Ribosome Affinity Purification followed by mRNA-seq) to RNA-seq. Wilcoxon rank-sum test was carried out for statistical analyses. **c**, RNA-seq (upper) and TRAP-seq (lower) data generated from the rice calli showing *Tos17* retrotransposon and its neighbouring gene (LOC_Os07g44710). **d**, Codon translational coefficient (CTC) plotted from the highest to the lowest. The optimal and suboptimal codons are colored in red and blue, respectively. Genes with at least FPKM 1 are only considered to calculate CTC. Inlets are the base compositions by codon nucleotide positions. nt, nucleotide.

Unequal usage of synonymous codons has been observed in many organisms and such codon sequence bias impacts on various RNA processes^43,44^. Codon optimality, a relative ratio of optimal to suboptimal codons, is often used as a proxy for certain RNA features of interest. Such codon index derived from sequence information is particularly useful in the study of translation because weakly expressing TE transcripts are less represented in the ribosome-associated fraction and thus difficult to be assessed precisely for their translational activities. In order to assign a general measure indicative to translational potential, we first categorized the rice codons by their correlativeness to translation (Fig. 1d). For this, we analysed the codon frequencies of the annotated transcriptional units of the rice genome. The Pearson’s correlation coefficient of codon frequency and TEI was defined as the codon translational coefficient (CTC) and each codon was assigned for its CTC (Fig. 1d). The expressed genes are only considered in CTC calculation to ensure that codon frequency well reflects the translational activity. As shown in Fig. 1d, the codons of rice exhibited varying levels of CTC. Noticeably, the codons ending with G or C showed positive CTC values meaning that those codons are more frequently used in the actively translating RNAs (Supplementary Fig. S1). On the other hand, A or U-ending codons showed low CTC values (Fig. 1d and Supplementary Fig. S1), which is reminiscent of the low GC3 of transposons shown in Fig. 1a. Taken together, transposons in the rice genome exhibit weak translation that is likely attributed to their codon sequence bias and the reduced translational activity may induce RNA cleavage presumably through the NGD pathway.

### Codon optimality negatively correlates with easiRNA production

Based on the CTC data obtained from Fig. 1, we defined those above 0.15 of CTC values as optimal codons and those below −0.15 as suboptimal codons (Fig. 1d). We then calculated the relative log2-ratio of optimal to suboptimal codon frequencies of each transcripts that will be hereinafter referred to as codon optimality. The codon optimality showed positive correlation with TEIs (Fig. 2a), while the out-of-frame false codon optimality showed less and insignificant correlation (Fig. 2b and c). This indicates that codon optimality well reflects the translatability. It is worth mentioning that although CTC was determined only from the expressed genes, the resulting codon optimality can be assigned to any transcriptional units including transposons as long as the coding sequence is provided.

**Figure 2.**
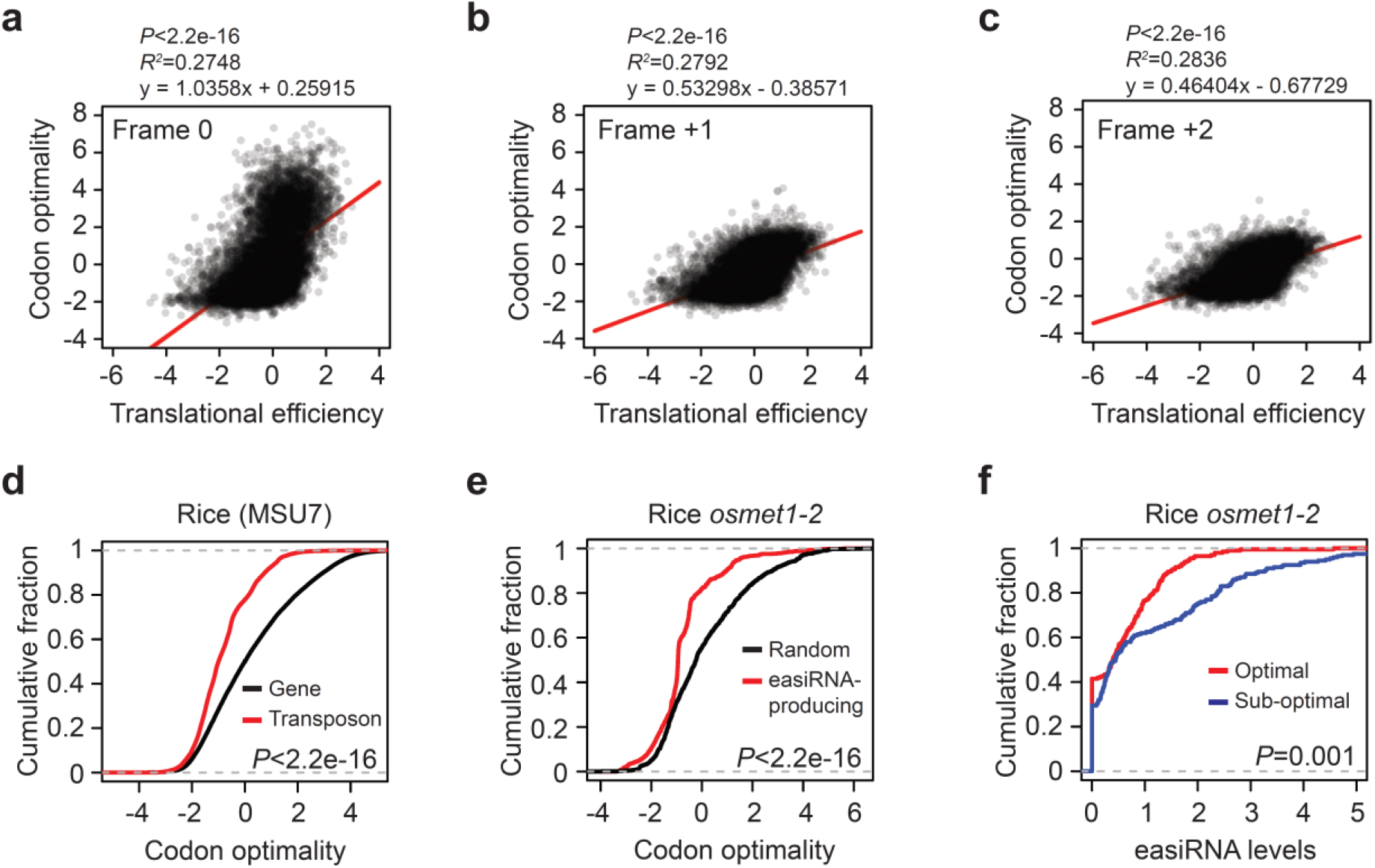
Codon optimality and translational efficiency. **a-c**, Correlation of codon optimality and TEI. Codon optimality was calculated from the codon sequences determined for the in-frame (**a**), frame +1 (**b**) and frame +2 (**c**). The entire transcriptional units annotated in the rice MSU7 genome containing both genes and transposons are plotted. Pearson’s product-moment correlation was used for statistical analyses (**a-c**). **d** and **e**, Comparisons of codon optimality between genes and transposons of rice (**d**) and the easiRNA-producing loci and randomly selected loci (**e**). The easiRNA-producing loci are the top 1,000 genes ranked by the easiRNA levels. **f**, The easiRNA levels in the suboptimal and optimal TEs in the rice *osmet1-2* mutant. The optimal and suboptimal TEs are the top 1,000 TEs when ranked from the highest and lowest codon optimality, respectively. The levels of easiRNAs were expressed as log2(FPKM+1). Wilcoxon rank-sum test was used for statistical analyses (**d-f**).

The codon optimality was then directly compared in genes and transposons of the rice genome. Figure 2d shows that transposons are significantly lower in the codon optimality, resembling the reduced translational activity shown in Fig. 1b. In order to see if the reduced translational activity and the low codon optimality of transposons is conserved in other species, we carried out ribosome footprint profiling sequencing (ribo-seq) experiments using the *decrease in dna methylation 1 (ddm1*) mutant of *Arabidopsis*. Similar to those of rice, *Arabidopsis* transposons were drastically reduced in translation and lower in codon optimality compared to genes (Supplementary Fig. 2), indicating that the codon sequence bias and translational repression of transposons is conserved in both monocot and dicot plants.

We next selected for the loci generating the easiRNAs in the rice *osmet1-2* mutant, defective in CG methylation^45^, and compared their codon optimality with randomly selected loci. As can be seen in Fig. 2e, the easiRNA-producing loci are lower in codon optimality, likely exhibiting weaker translational activities. Oppositely, the optimal and suboptimal transposons were selected by the codon optimality and compared for their easiRNA levels. Consistently, the suboptimal TEs produced higher levels of easiRNAs (Fig. 2f). In summary, the suboptimal codon usage and the reduced translational activity of TEs is conserved in plants and correlates with active easiRNA production.

### Ribosome stalling triggers RNA cleavage

Translational inhibition causes ribosome stalling and in severe cases ribosome stacking or queuing^40,46,47^. In order to test if the reduced translational activity of transposons induces the RNA truncation which is a critical requirement for RDR6 targeting, we investigated the degradome-seq data generated from *ddm1* mutant of *Arabidopsis*^8^ Degradome-seq technique sequences the 5’ end of the truncated RNAs and from this we determined the degradability by normalizing its levels by the RNA-seq levels. Shown in Fig. 3a is the degradability of high and low TEI genes, revealing that lowly translating mRNAs are more prone to truncation (Fig. 3b). Since the siRNAs can inhibit the translational activity of mRNAs^48^, we wanted to test whether the weak translation of transposons is the cause or consequence of siRNA function. For this, we carried out additional ribo-seq experiments using the *ddm1 rdr6* double mutant which does express transposons but does not produce easiRNAs. Interestingly, we were not able to detect any noticeable changes of translational activities of transposons between *ddm1* and *ddm1 rdr6* double mutants (Fig. 3c). The RDR6 target TEs (Fig. 3c, dots marked in red) also exhibited comparable levels of translational efficiency in both mutants, indicating that the reduced translation of TEs is less likely caused by siRNAs but rather by unfavourable codon sequence usage.

**Figure 3.**
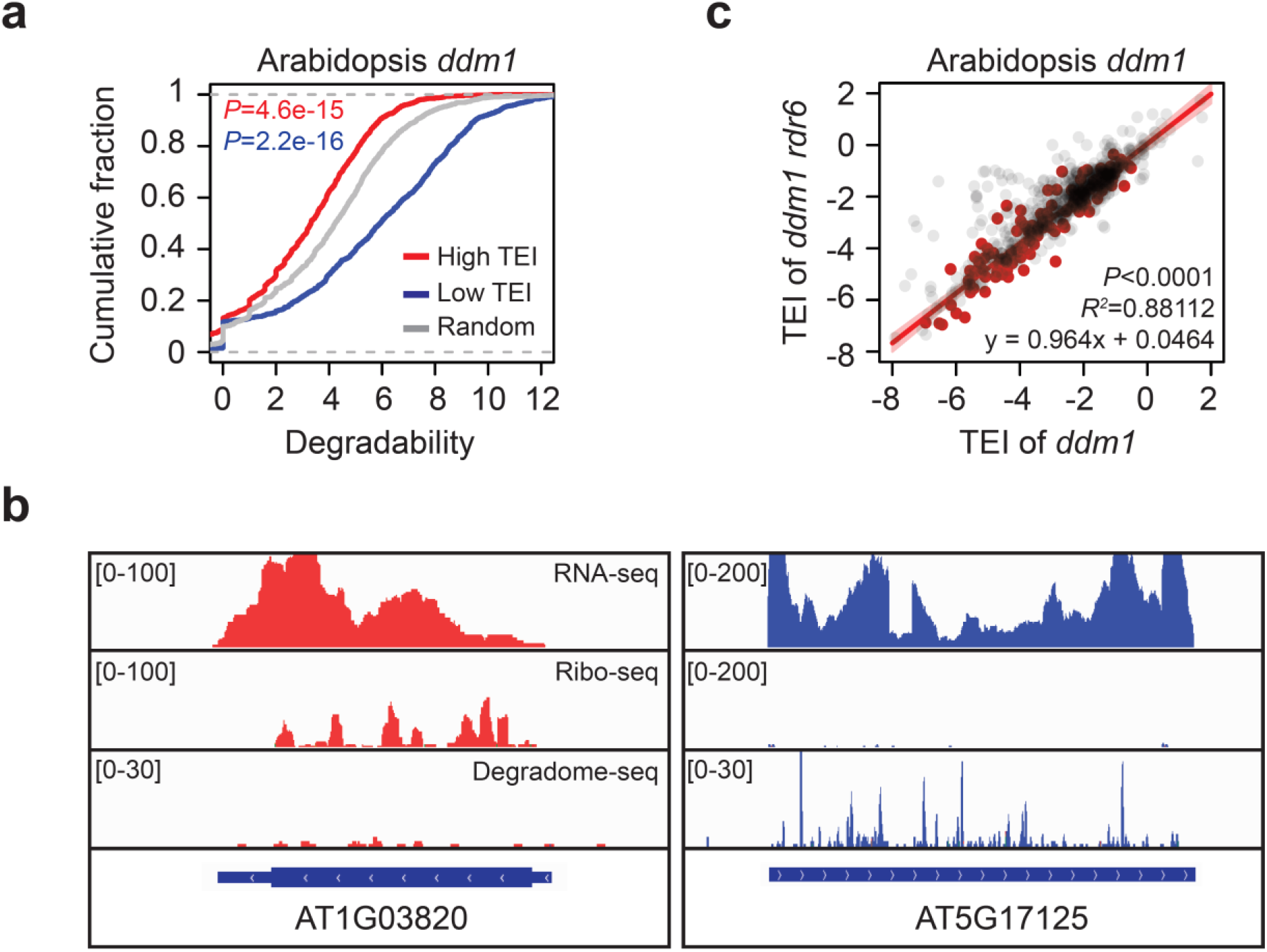
Translational inhibition causes RNA truncation. **a**, Degradability of high TEI, low TEI and random genes in *ddm1* mutant of *Arabidopsis.* The high and low TEI genes are the top 1,000 genes when ranked from the highest and lowest TEI genes, respectively. Degradability was determined by log2-fold change of degradome-seq normalized to RNA-seq. *P* values were obtained for the high TEI (red) and low TEI genes (blue) compared to the random genes by Wilcoxon rank sum test. **b**, Representative loci for actively translating (red) and weakly translating genes (blue) in *ddm1* mutant. From top, RNA-seq, Ribo-seq and degradome-seq. **c**, Translation efficiency of *ddm1* and *ddm1 rdr6* double mutant. Red dots represent RDR6 targets identified for those with reduced easiRNA levels in *ddm1 rdr6* double mutants than in *ddm1* by the log2-fold change below −1. Grey dots are expressed transposons with FPKM larger than 1. Pearson’s product-moment correlation was used for statistical analyses.

A previous study showed that collision of stacked ribosomes is critical for triggering NGD pathway^37^. Given this, we reasoned that transcripts containing the stacked ribosomes might be more frequently truncated and thus readily processed to easiRNAs. In order to profile the RNAs containing the queued ribosomes, we selected from *ddm1* ribo-seq data the di-ribosome (disome) fragment reads ranging from 40 to 65 bp (Fig. 4a). Disome fragments were strongly enriched with the non-protein-coding RNAs including tRNAs and rRNAs as well as organellar RNAs, while only around 20 % was protein-coding genes (Fig. 4b), which is overall consistent with the previous reports^40,47^. Noticeably, disome RNAs are more strongly enriched with TEs (Fig. 4c), further supporting the notion that TE transcripts are associated with translational repression. To analyse the coding sequence features of disome RNAs, we retrieved the sequences of the disome-containing protein-coding genes and compared the codon optimality with randomly selected RNAs. As shown in Fig. 4d and e, disome RNAs showed drastically reduced codon optimality and translational efficiency, suggesting that ribosome stalling might be caused by the codon sequence usage unfavourable for translation. More importantly, disome RNAs are significantly more prone to RNA cleavage than randomly selected RNAs (Fig. 4f). We then determined the easiRNA levels of disome RNAs and indeed they produced considerably more easiRNAs than randomly chosen RNAs (Supplementary Fig. S3a). Oppositely, RDR6 target transposons were selected according to the dependency of easiRNA production on RDR6 and their codon optimality was measured. Consistently, we observed that RDR6 targets have lower codon optimality compared with randomly selected non-RDR6 targets (Supplementary Fig. S3b). In conclusion, ribosome stalling resulted from the suboptimal codon usage triggers RNA cleavage and subsequently easiRNA production.

**Figure 4.**
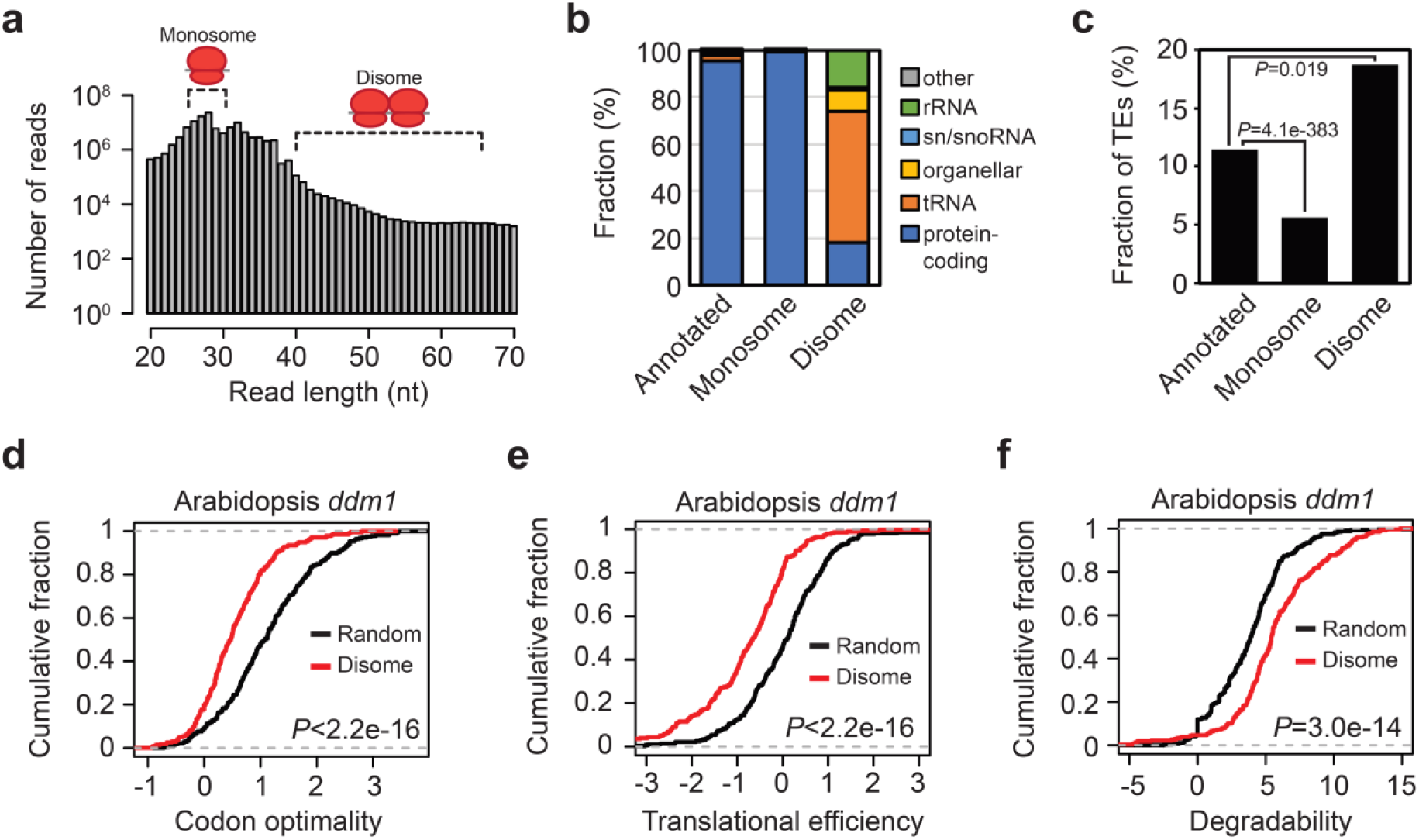
Ribosome queuing and RNA truncation. **a**, Numbers of reads by read lengths of ribo-seq generated from *ddm1*. Reads ranging from 40 to 65 bp were selected as disome fraction. **b**, Genomic features of disome loci compared with the entire annotated genes and monosome loci of *Arabidopsis.* Monosome and disome loci were selected for those above 1 of FPKM. **c**, The fraction of transposons in the annotated, monosome- and disome-containing genes. Hypergeometric test was performed to obtain *P* values. **d-g**, Comparison of disome RNAs for codon optimality (**d**), translation efficiency (**e**) and degradability (**f**). Data generated from *ddm1* mutant was used for the analyses (**d-f**). Wilcoxon rank-sum test was carried out for statistical analyses (**d-f**).

### Liquid-liquid phase separation mediates the formation of siRNA bodies and SGs

So far, we have shown that the unique codon sequence usage of transposons led to ribosome stalling and thereby RNA truncation which is an important prerequisite for the easiRNA production. Apart from the RNA truncation, the specificity of the easiRNA pathway is given by the spatial isolation of cytoplasmic compartments known as siRNA bodies^49–52^. It is well documented that non-membranous cellular compartments are formed by the liquidliquid phase separation of ribonucleoproteins^27^. Prion-like domains or low complexity sequences are common protein features that are frequently associated with phase-separating proteins^53^. Although RDR6 and SGS3 form the cytoplasmic siRNA bodies and SGS3 contains the prion-like domains (Fig. 5a), their biophysical property of LLPS has never been investigated so far. In order to test if SGS3 indeed undergoes LLPS, the GFP-tagged SGS3 protein of *Arabidopsis* was expressed in *E. coli* and purified for the *in vitro* phase separation assay. As shown in Fig. 5b, while the GFP alone does not form any globular protein condensates, the full-length SGS3 protein forms the liquid droplets *in vitro* which is a typical feature of LLPS. In order to demonstrate that the phase separation behaviour of SGS3 is dependent on the prion-like domains, we deleted the prion-like domains and purified the truncated SGS3 proteins. The SGS3 protein without prion-like domains did not exhibit any phase-separating behaviour, suggesting that the LLPS of SGS3 is mediated by the prion-like domains (Fig. 5b). To further demonstrate the fluidity and dynamicity of SGS3 protein droplets, which is an important characteristic of phase-separating proteins, we carried out the time-lapse microscope imaging analysis of SGS3 protein droplets. Figure 5c shows the fluorescence microscope images of two adjacent SGS3 protein droplets which are fusing together within only several seconds (Supplementary Movie S1). Additionally, we performed Fluorescence Recovery After Photobleaching (FRAP) assay and observed that the lesions of the photobleached SGS3 protein droplets recovered almost completely in around 30 seconds (Fig. 5d and e; Supplementary Movie S2). These altogether indicate that LLPS is an important physiochemical feature of SGS3 acting as a critical driving force for the siRNA body formation.

**Figure 5.**
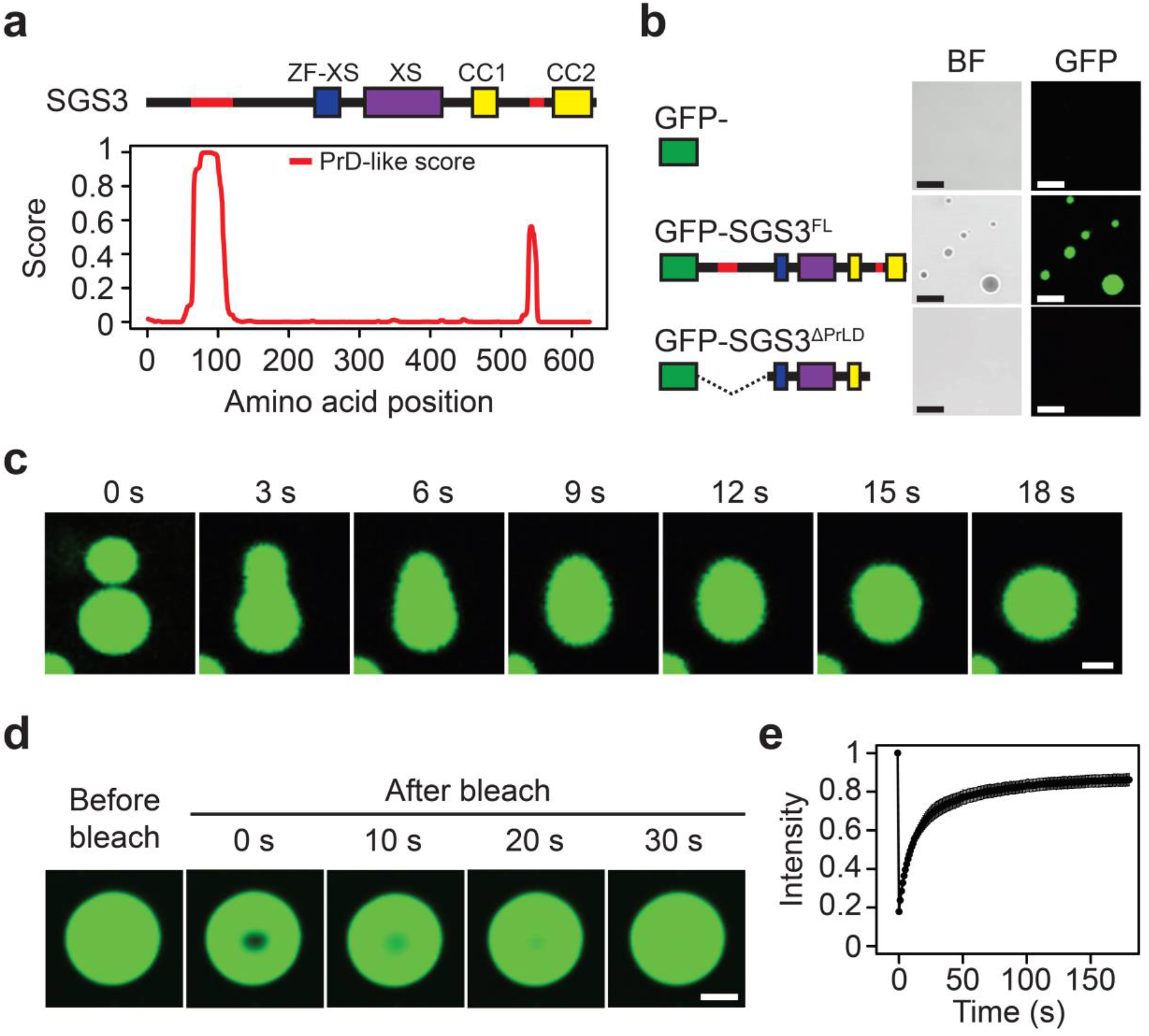
*In vitro* phase separation assay of SGS3. **a**, Protein domain structure (upper) and PrD-like score (lower) of SGS3. ZF-XS, zinc finger-rice gene X and SGS3; CC, coiled coil; PrD, prion-like domain. **b**, Bright field (left) and fluorescence (right) microscope images of GFP, GFP-tagged SGS3 full length protein and GFP-tagged SGS3 protein without prion-like domains. BF, bright field; FL, full length; PrLD, prion-like domain. Scale bars, 10 μm. **c**, Fluorescence time-lapse microscope images of GFP-SGS3^FL^. Scale bar, 2 μm. **d** and **e**, FRAP of GFP-SGS3^FL^ shown as the time-lapse fluorescence microscope images (**d**) and the plot for the fluorescence intensity during the time course of recovery after photobleaching (**e**). Scale bar, 5 μm. Data are presented as mean ± s.d. (*n* = 13).

It has been well documented that the siRNA body components colocalize with UBP1b, a major stress granule (SG) core component^48,49^. Studies in human cells revealed that the formation of SGs is mediated by the LLPS of TIA-1, a homolog of UBP1b, through its prion-like domains^29,30^. The SGs in plants are formed when plant cells are stressed and in the DNA methylation-deficient *ddm1* mutant of *Arabidopsis^22,48^*. Unfortunately, the plant UBP1b protein has never been assessed so far for its phase separation behaviour. The *Arabidopsis* UBP1b contains two prion-like domains at its both ends (Supplementary Fig. S4a), potentiating its prion-like behaviour. Indeed, our *in vitro* phase separation assay revealed that UBP1b undergoes LLPS and the prion-like domains are required for the phase separation behaviour (Supplementary Fig. S4b). Intriguingly, the phase separation activity of UBP1b was reduced as *Arabidopsis* RNAs are supplemented in the assay buffer (Supplementary Fig. S4c). This is in agreement with a previous study that RNA inhibits phase separation behaviour of prion-like RNA-binding proteins to prevent the aberrant formation of protein condensates^54,55^. Taken together, the formation of plant SGs and their accompanying siRNA bodies is mediated by LLPS of UBP1b and SGS3, respectively.

### Weakly translating RNAs are preferentially guided to cytoplasmic foci

While the protein components of SGs and siRNA bodies in plants are relatively well known^56^, the RNA composition of such cytoplasmic compartments are poorly understood. Studies in human and yeast have shown that transcriptome of cytoplasmic RNA granules are associated with strong translational repression^23,25,57,58^. This led us to hypothesize that transposon RNAs might be preferably located to cytoplasmic foci including SGs and siRNA bodies presumably owing to their weak translational activities and are therefore selectively taken over to the RDR6-RdDM pathway. In order to demonstrate this hypothesis, we enriched the cytoplasmic body fraction of *ddm1* mutant using the previously established SG enrichment method^25,59^ and sequenced the RNAs (SG-RNA-seq). By normalizing the SG-RNA-seq levels to the total RNA-seq levels we assessed the SG-enrichment of each transcript and identified 863 SG-enriched and 891 SG-depleted RNAs (Fig. 6a). Noticeably, the fraction of transposons in the SG-enriched RNAs was over 35 %, while those of the SG-depleted RNAs and the expressed transcripts were only around 5 % (Fig. 6b). This data strongly supports the notion that TE transcripts in *Arabidopsis* are strongly enriched in the cytoplasmic foci. Similarly, a previous study in human cells suggested that AU-rich transcripts are strongly enriched in the cytoplasmic RNA granules^60^, resembling the sequence feature and cellular localization behaviour of transposons in plants. Our SG-RNA-seq also revealed that SG-enriched RNAs are remarkably lower in the RNA levels (Fig. 6c), codon optimality (Fig. 6d) and translational efficiency (Fig. 6e) but associated with higher levels of easiRNAs (Fig. 6f and g). We then directly compared the SG-enriched TEs and the targets of RDR6 and DDM1 of *Arabidopsis,* which revealed a substantially large proportion of overlapping transcripts (Supplementary Fig. S5). These data collectively indicate that TE RNAs are preferentially localized to cytoplasmic compartments including SGs and siRNA bodies where the easiRNA pathway is present, providing additional selectivity of RDR6-RdDM towards transposon RNAs.

**Figure 6.**
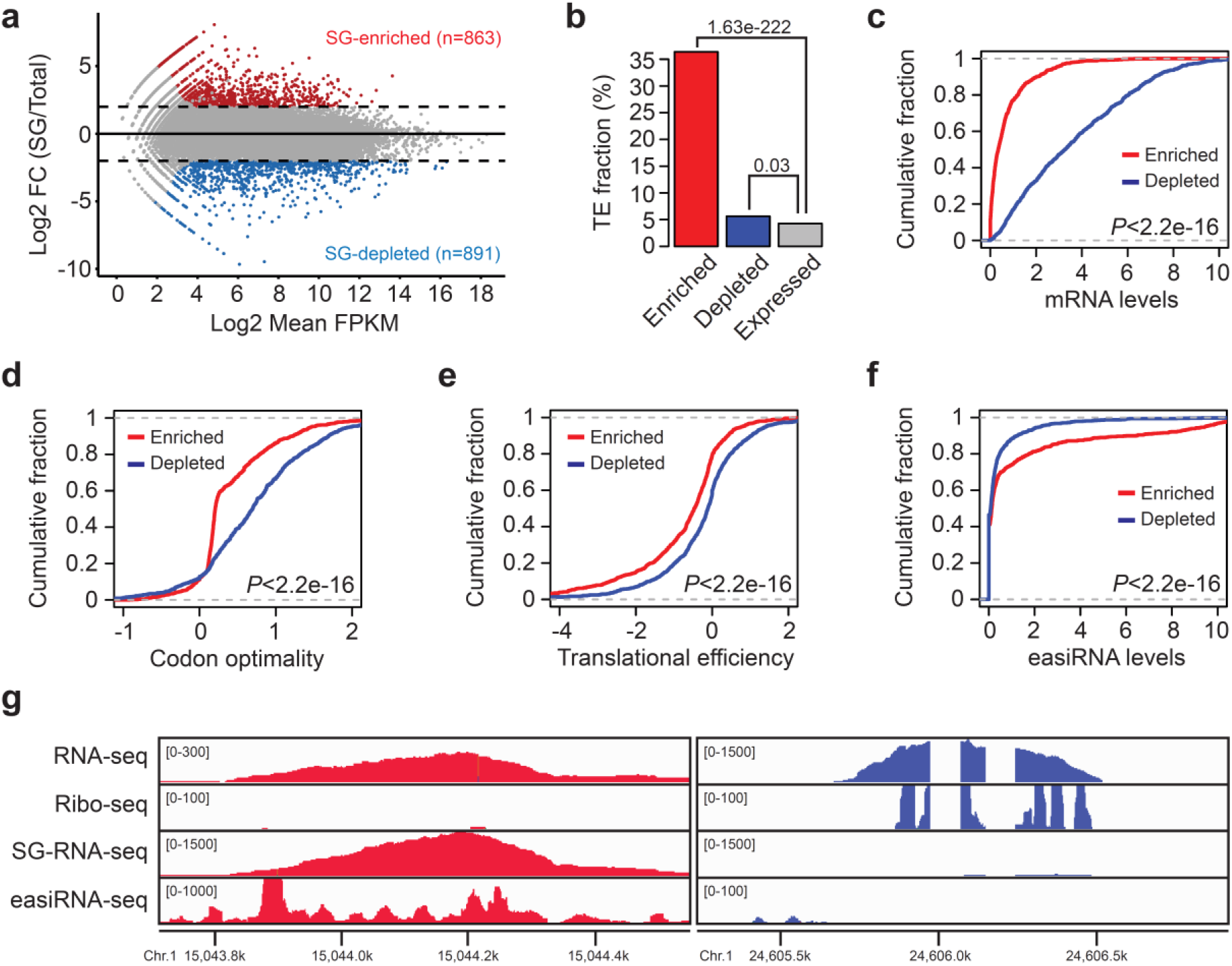
Localization of transposon RNAs to cytoplasmic foci. **a**, MA plot for the RNA-seq of the stress granule (SG)-containing cytoplasmic compartments (SG-RNA-seq). SG-enriched (red) and depleted (blue) transcripts are identified by the log2-fold change hgher than 2 or lower than −2 and FDR values below 0.05. **b**, Fraction of transposons in SG-enriched and depleted transcripts. Genes with FPKM value above 1 were defined as being expressed. Hypergeometric test was used to obtain *P* values. **c-f**, Comparison of SG-enriched and depleted RNAs for the mRNA levels (**c**), codon optimality (**d**), translation efficiency (**e**) and easiRNA levels (**f**). The levels of easiRNAs are expressed as log2(FPKM+1). Wilcoxon rank-sum test was performed for statistical analyses (**c-f**). **g**, Genomic loci showing the RNA-seq, ribo-seq, SG-RNA-seq and easiRNA-seq of SG-enriched (left) and depleted (right) TE.

## Discussion

RDR6-RdDM is a critical cellular pathway in plants that detects and suppresses transposon RNAs. The aberrancy of mRNAs derived from genes are mitigated predominantly by RNA decay pathways and RDR6-RdDM is usually prevented because the resulting siRNAs may target the normal transcripts^18,61–64^. As illustrated in Fig. 7, we suggest that transposon RNAs, unlike genic mRNAs, are specifically detected and targeted to RDR6-RdDM by their reduced translational activities. The weak translation of TE RNAs contributes to both RNA truncation and localization to siRNA bodies which are both important for the selective easiRNA processing of transposon RNAs. Although RDR6 can target several endogenous genes to produce *trans*-acting siRNAs in normal condition^13,22,65^, the major RNA templates of RDR6 are those from foreign and invasive genetic elements such as virus, transgenes and transposons^11,33,66^. Given that non-self or alien RNAs are presumably less optimal to the host’s codon usage and commonly regulated by the similar siRNA-mediated pathway, the epigenetic silencing of viral RNAs and transgenes might be triggered by the translation-coupled pathway as was seen in transposons. Therefore, our work can provide a novel framework for treating plant viral diseases and improving genetic engineering through transgene transformation.

**Figure 7.**
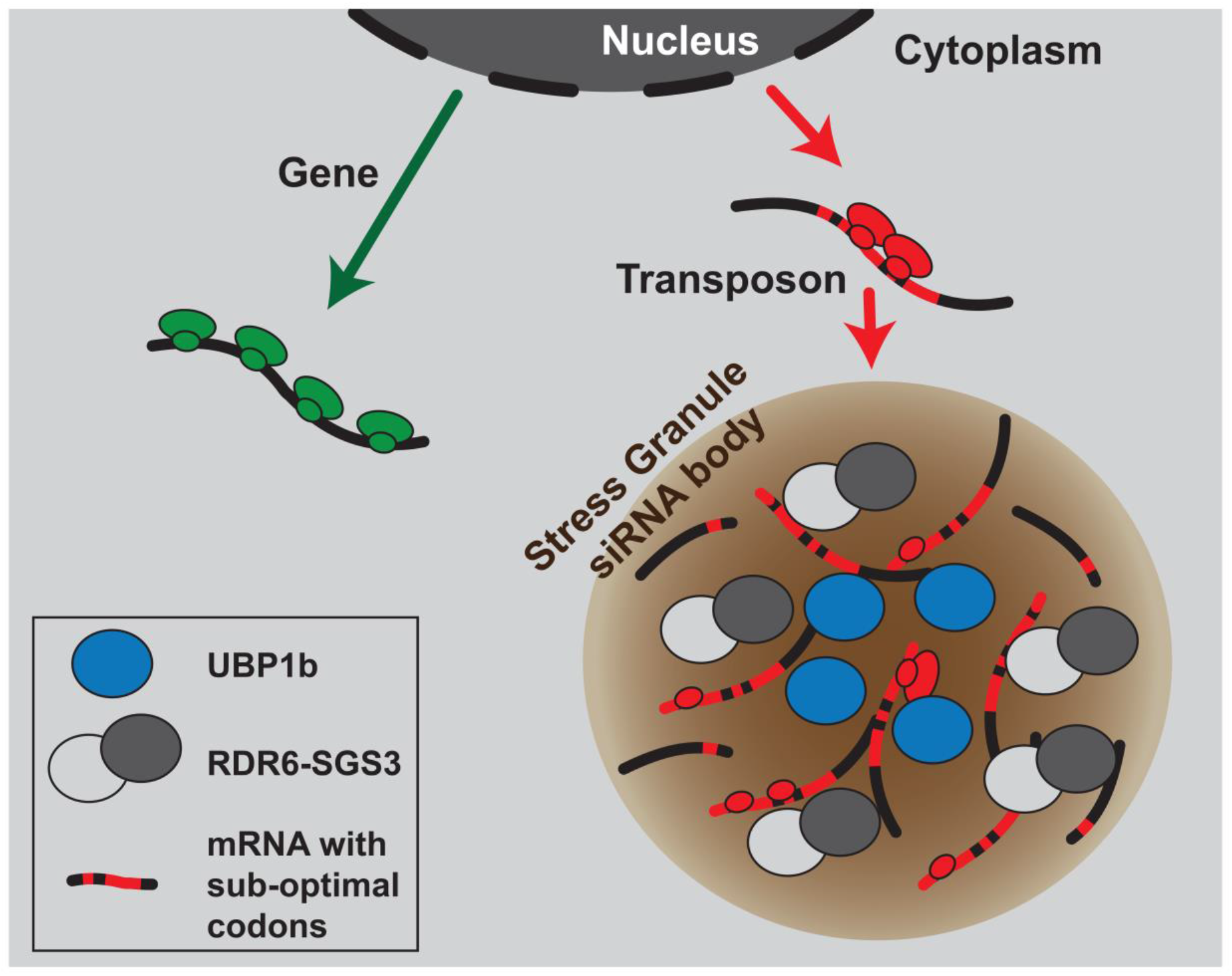
A schematic model for the specific recognition of transposon RNAs and the initiation of easiRNA biogenesis. Unlike genic mRNAs, transposon RNAs exhibit reduced translational efficiency because of the suboptimal codon usage. Ribosome stalling leads to RNA truncation and localization to cytoplasmic foci which collectively contribute to selective processing of transposon RNAs to easiRNAs.

We have demonstrated that the suboptimal codon usage and reduced translation of transposons is conserved in rice and *Arabidopsis.* Several other studies also showed that localization to SGs (where mRNA translation is inhibited) and ribosome stalling frequently occurs to human retroelements such as LINE-1 and *Alu*^67–69^. Our investigation to GC contents in several genomes of animals and invertebrates also revealed that GC3 is drastically reduced in transposons (Supplementary Fig. S6), partly indicating that the weak translation might be a general feature of transposons in eukaryotes. In addition, it has been suggested that stalled spliceosome caused by suboptimal splice sites triggers RNAi in yeast^70^. These altogether may suggest an interesting notion that certain abnormalities of RNA processing might have been selected to serve as a signal to turn on the genome surveillance system of the hosts. Although we have shown in this study that the ribosome-stalled TE RNAs are preferentially funnelled to the easiRNA production pathway in the siRNA bodies, the precise regulatory mechanisms for the specific guidance to particular cytoplasmic compartments are yet to be concluded. Moreover, the NGD pathway was suggested to trigger RNA truncation at the ribosome-stalled sites, however, whether the RDR6-RdDM requires the NGD or not is to be confirmed.

RDR6 and AGO7 are known to function in conjunction with SGS3 and colocalize together in the cytoplasmic siRNA bodies^49^. However, we were not able to find any prionlike domains in RDR6 and AGO7 (Supplementary Fig. S7), suggesting that they might be guided to siRNA bodies possibly via the physical interaction with SGS3. Several other studies have shown that LLPS can be enhanced when additional biomolecules are supplemented^55,71,72^. In this regard, the functional role of the interacting proteins on the phase separation behaviour of SGS3 and the formation of siRNA bodies will be worth to be followed and investigated. In addition, our interrogation of prion-like domains in the small RNA pathway factors revealed that AGO1, 2, 3 and 5 contain prion-like domains at their N termini (Supplementary Fig. S7). This may suggest that apart from the easiRNA pathway, other cellular processes involving small RNAs in plants can also be mediated by LLPS.

In summary, the specific recognition of transposons and their targeting to siRNA biogenesis pathway is critical for perpetual maintenance of the genome integrity. In plants, the host genomes detect and discriminate transposon RNAs by their reduced translational activity. The weak translation then causes RNA truncation and siRNA body localization which together provide selectivity to TE RNAs for siRNA production. The specificity of siRNA pathway is further secured by the formation of siRNA body which is mediated by the phase separation of SGS3 protein.

## Methods

### Plant materials and growth condition

*Arabidopsis* seeds of Columbia-0 (Col-0), *ddm1-2* (selfed for five generations) and *ddm1-2 rdr6-11* double mutants (genotyped from F2 segregation population derived from *ddm1-2* and *rdr6-11* crosses) were surface-sterilized in 75 % ethanol and germinated on half-strength Murashige and Skoog media. Plants grown for 10 days under 16 h light/8 h dark cycling at 22 °C were collected for RNA-seq, SG-RNA-seq and ribo-seq.

### Codon sequence analysis

Codon frequency was calculated for the coding sequences of rice and *Arabidopsis* using the R package “seqinr”. We used rice MSU7 and *Arabidopsis* TAIR10 version of genome assembly and annotation downloaded from http://rice.plantbiology.msu.edu/pub/data/Eukaryotic_Projects/o_sativa/annotation_dbs/ for rice and ftp://ftp.arabidopsis.org/home/tair for *Arabidopsis.* Codon translation coefficient was defined as the Pearson’s correlation coefficient between the codon frequency and translation efficiency of genes that have the FPKM value of at least 1. Optimal codons and suboptimal codons (Fig. 1d) are those above 0.15 and below −0.15 of CTC, respectively. The log2-ratio of optimal to suboptimal codon frequency of individual transcript is defined as codon optimality.

### Next-generation sequencing (NGS) library construction

For RNA-seq, the mRNAs were purified from 3 μg of total RNA using poly-T oligo-attached magnetic beads. Library preparation was carried out using the NEBNext^®^ UltraTM RNA Library Prep Kit (NEB) following the manufacturer’s instruction. Sequencing was performed on an Illumina HiSeq platform and 150 bp paired-end reads were generated.

For ribo-seq, the plant samples were lysed and digested by RNase I, then ribosome protected fragments (RPFs) were purified using the MicroSpin S-400 columns (GE Healthcare). After rRNA depletion, the RPFs were purified by polyacrylamide gel electrophoresis (PAGE). Then, the 5’ and 3’ adapters were ligated following the end-repair and dA-tailing. The adapter-ligated cDNAs were obtained by the one-step reverse transcription and PAGE purification. After PCR amplification and PAGE purification, the sequencing library was prepared using the NEBNext^®^ Multiplex Small RNA Library Prep Kit (NEB) and the resulting library was loaded onto an Illumina HiSeq X machine for PE150 sequencing.

### NGS data analysis

For RNA-seq data analysis, the raw data were first processed through the in-house Perl scripts to remove reads containing adapter, ploy-N and low-quality sequences. Clean reads were then aligned to the rice (MSU7) and the *Arabidopsis* reference genome (TAIR10) in default settings using Hisat2 (version 2.0.5). The FPKM of gene and transposons were calculated by StringTie (version 1.3.5). Visualization of the sequencing data was performed using the Integrative Genomics Viewer (IGV).

For ribo-seq data analysis, the software Cutadapt (version 1.12) was first used to trim adapter sequences and the reads between 20-50 bp were retained. FASTX_toolkit (version 0.0.14) was used to filter out the low-quality reads and Bowtie (version1.0.1, parameter –l 20) to filter out the structural and ribosomal RNA reads. The kept reads were aligned to the genome by Tophat2 and the cufflinks (version 2.2.1) were employed to calculate FPKM. For disome analysis, the reads between 40-65 bp after removal of adapters were selected and no filtering was performed for the non-coding RNAs.

Public datasets used in this study are from PRJNA298638 (rice TRAP-seq)^35^, SRP043448 (rice small RNA-seq)^45^ and GSE52952 *(Arabidopsis* small RNA-seq and degradome-seq)^8^.

### Enrichment of cytoplasmic bodies

The enrichment of cytoplasmic bodies was carried out by employing the SG enrichment methods reported previously^25,56,59^. Briefly, 2 g of samples was ground with a precooled mortar and pestle in liquid nitrogen. The ground samples were collected into 50 ml conical tube and resuspended in 5 mL of SG lysis buffer (50 mM Tris-HCl pH 7.4, 100 mM KOAc, 2 mM MgOAc, 0.5 mM DTT, 0.5% NP40, Complete EDTA-free Protease Inhibitor Cocktail (Roche), 1 U/mL of RNasin Plus RNase Inhibitor (Promega)). The resulting slurry was centrifuged at 4,000 g for 10 min at 4 °C, the supernatant was removed, and the pellet was resuspended in 2 ml of lysis buffer. The samples were again centrifuged at 18,000 g for 10 min at 4°C. The pellets were resuspended in 2 ml lysis buffer, vortexed and centrifuged at 18,000 g at 4 °C for 10 min. The supernatant was discarded and the pellets were resuspended in 1 ml of lysis buffer. After a final centrifugation at 850 g for 10 min at 4°C, the supernatant (enriched with SGs) was transferred to a new 1.5 ml microcentrifuge tube and purified using the RNeasy Plant Mini Kit (QIAGEN).

### Protein sequence analysis

Protein domains were predicted by SMART (http://smart.embl-heidelberg.de/) using the full-length amino acid sequences. Prediction of prion-like domains was performed using the web-based tool, PLAAC (http://plaac.wi.mit.edu/).

### Protein expression and purification

To produce the recombinant proteins, the coding sequences of *Arabidopsis* genes were PCR amplified from the reverse-transcribed cDNAs prepared from Col-0 seedling samples using the specific primers listed in the Supplementary Table S1. The amplified DNA was then cloned into the modified pET28a (Novagen) expression vector containing the N-terminal eGFP that was introduced between BamHI and EcoRI sites. The expression of GFP-SGS3 protein was induced in *Escherichia coli* Rosetta (DE3) (Novagen) by adding 0.1 mM isopropyl β-d-1-thiogalactopyranoside (IPTG) at 16 °C overnight. The collected cells are resuspended in the lysis buffer (20 mM Tris-HCl pH 7.6, 200 mM NaCl, 10 % Glycerol, 0.1 % Tween20, 1 mM PMSF) and lysed by sonication, then centrifuged at 20,000 g for 45 min at 4 °C. The supernatants were purified with Ni-NTA (Qiagen) in the elution buffer (250 mM imidazole in lysis buffer) according to the manufacturer’s instructions and further purified using the Superdex 200 increase 10/300 column. The purified proteins were stored in the storage buffer (20mM HEPES pH 7.4, 150 mM KCl, 1 mM DTT) at 100 μM of protein concentration until used.

### *In vitro* phase separation assay

For *in vitro* liquid droplet assembly, 10 μM of purified proteins mixed with PEG8000 (NEB) at 10% (w/v) was used. GFP fluorescence was imaged using a Zeiss LSM880 confocal microscopy equipped with 40×/1.1 water immersion objective and the GaAsP spectral detector. The GFP was excited at 488 nm and detected at 491-535 nm.

### Microscopy analysis

For time-lapse microscopy, GFP fluorescence was observed under Zeiss LSM880 confocal microscope. Images were acquired every 3 sec for 5 min. At each time point, maximum projections from z-stack of 14 steps with step size of 0.6 μm were applied. Image analysis was performed with FIJI/ImageJ.

FRAP assay of GFP-SGS3 was performed on a Zeiss LSM880 Airy scan confocal microscope. Photobleaching was done using a 488 nm laser pulse. Recovery was recorded every second for 5 min.

### Resource availability

The NGS data generated in this study are deposited to SRA repository under PRJNA598331 [https://www.ncbi.nlm.nih.gov/sra/PRJNA598331]. The analyses were performed using the standard codes instructed by the tools described in the Methods.

## Acknowledgments

We would like to thank Yunxiao He from the Core Facility Center, Shanghai Institute of Plant Physiology and Ecology, Chinese Academy of Sciences for the technical support on the confocal microscopy. This work was supported by the Strategic Priority Research Program of Chinese Academy of Sciences (XDB27030209) and the National Natural Science Foundation of China (31970518) granted to J.C.

## Author contributions

J.C. conceived the idea and designed the experiments. E.Y.K., Z.L., H.L. and W.F. conducted the experiments. L.W., E.Y.K., Z.L. and J.C. analysed the data, wrote and revised the manuscript.

---

**Supplementary Fig. S1.**
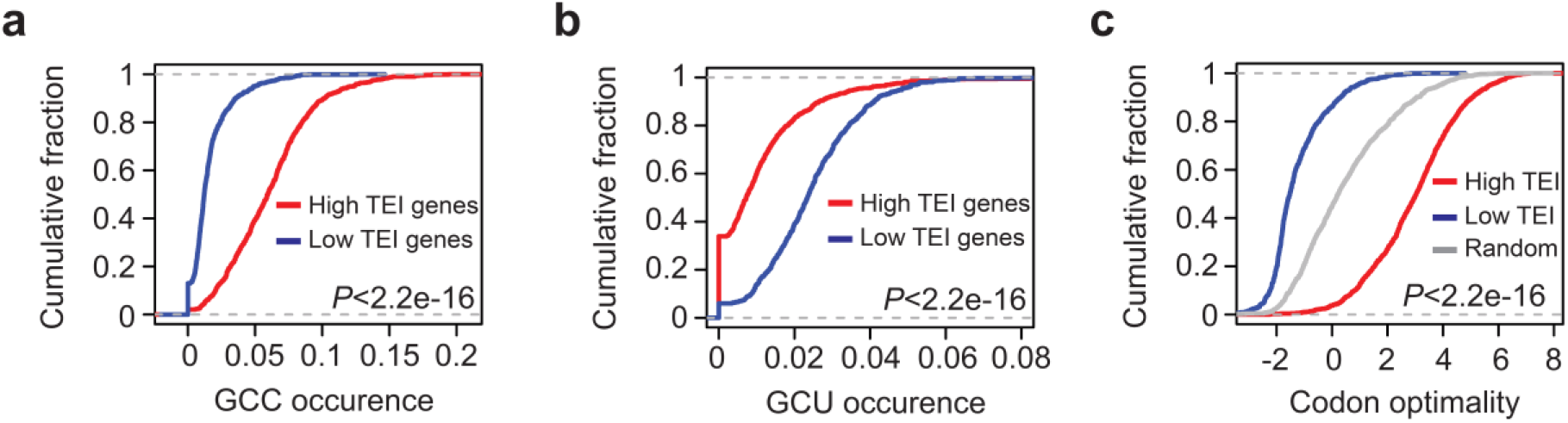
Codon frequency and translation efficiency in rice. **a** and **b**, GCC (**a**) and GCU (**b**) codon frequency in highly (red) and lowly translating mRNAs (blue). High and low TEI genes are top 1,000 genes when ranked from the highest and lowest TEI, respectively. TEI, translation efficiency index. **c**, Codon optimality of high TEI, low TEI and random genes. Wilcoxon rank-sum test was used for statistical analyses (**a-c**).

**Supplementary Fig. S2.**
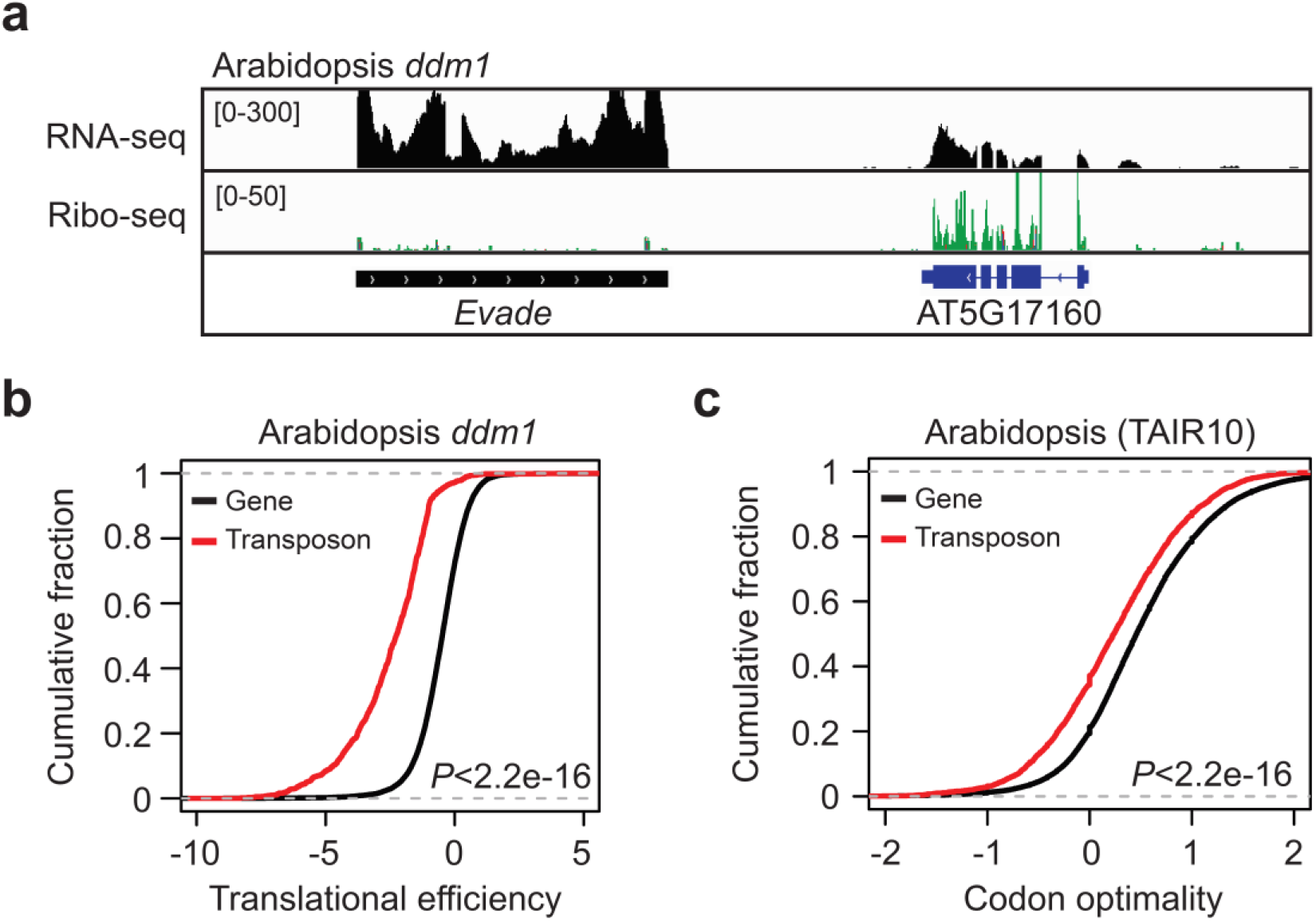
Translation efficiency and codon optimality of *Arabidopsis.* **a**, Genomic loci of *Arabidopsis* showing *Evade* retrotransposon and its neighbouring gene (AT5G17160) for RNA-seq (upper) and ribo-seq (lower). **b** and **c**, Comparison of genes and transposons in *Arabidopsis* for translation efficiency (**b**) and codon optimality (**c**). Translation efficiency and codon optimality was as determined in Fig. 1 and 2. Wilcoxon rank-sum test was carried out for statistical analyses (**b** and **c**).

**Supplementary Fig. S3.**
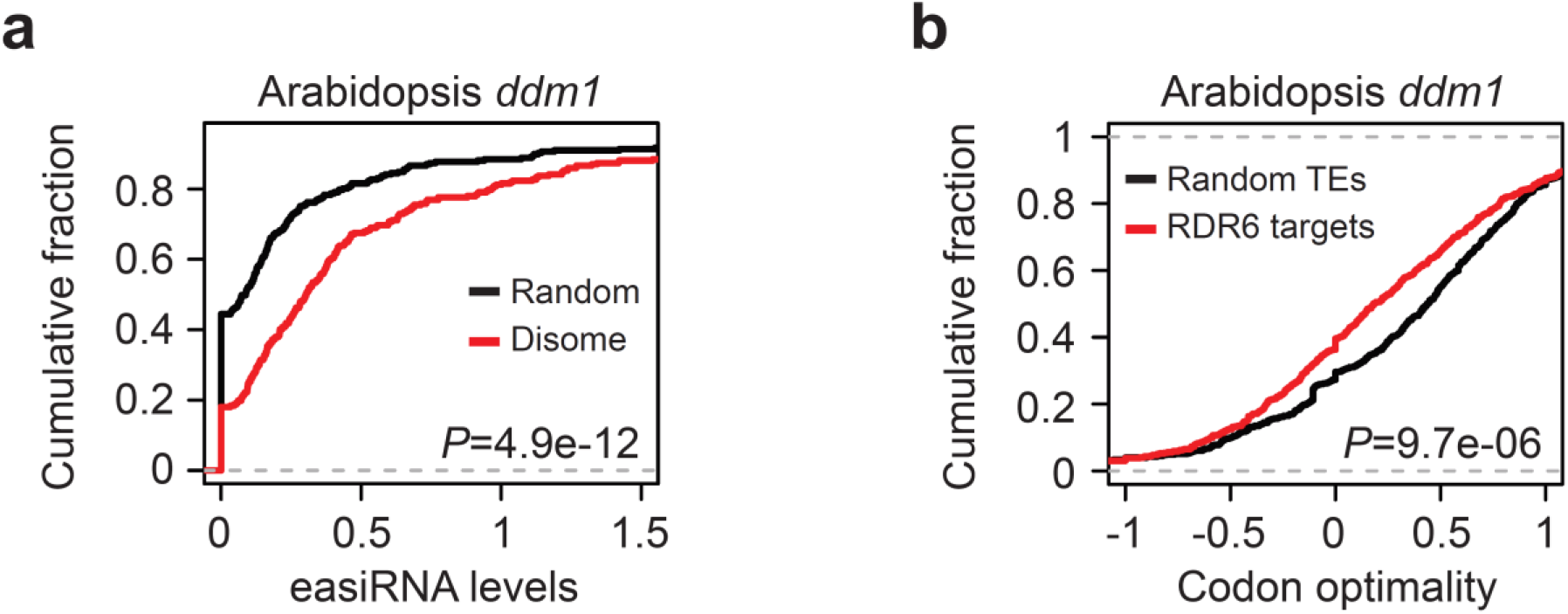
Ribosome stalling and easiRNA production. **a**, Comparison of disome RNAs for the easiRNA levels. The levels of easiRNAs are expressed as log2(FPKM+1). **b**, Codon optimality of RDR6 target transposons. RDR6 targets were identified as in Fig. 3c. Wilcoxon rank-sum test was carried out for statistical analyses.

**Supplementary Fig. S4.**
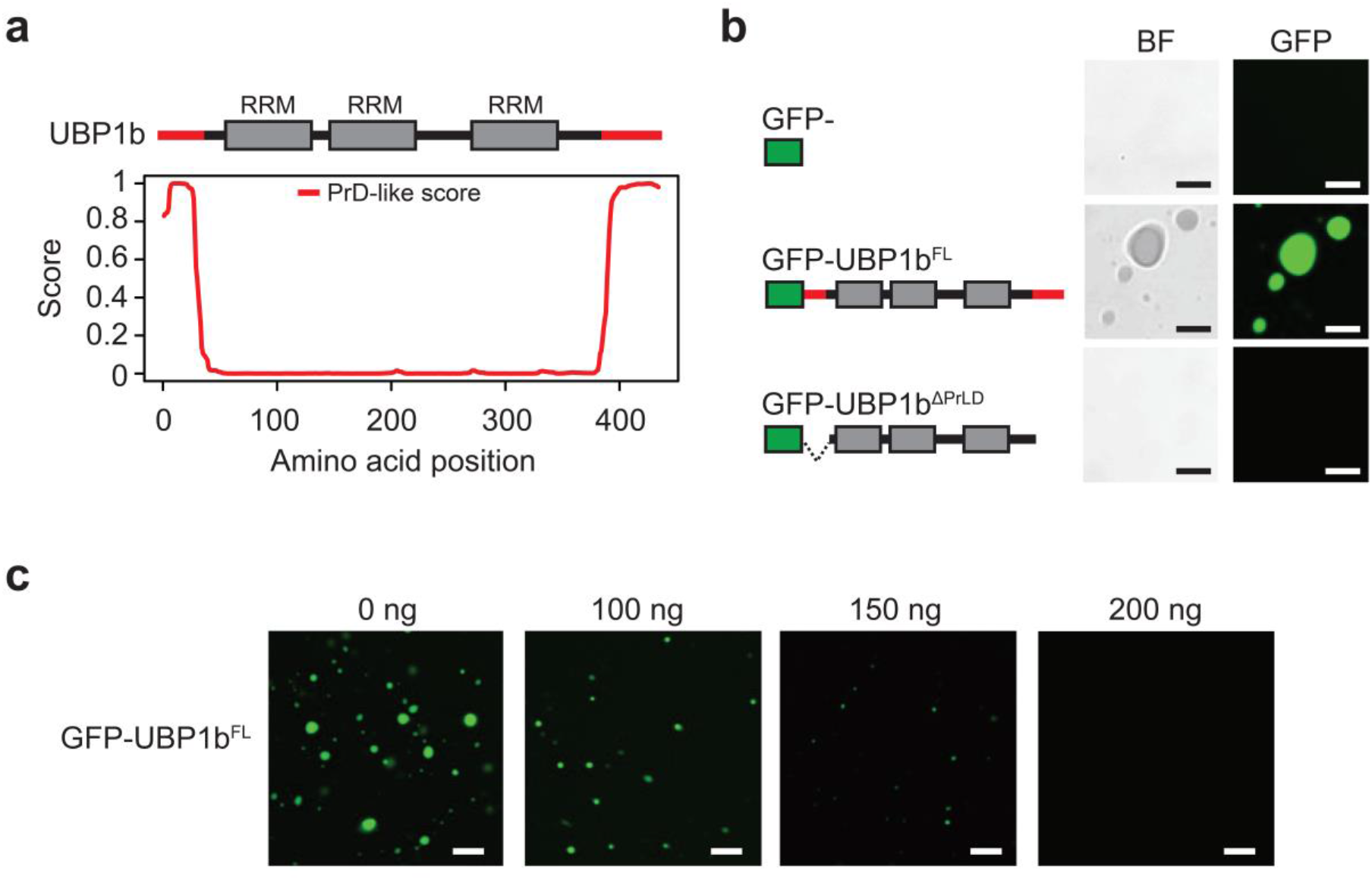
Liquid-liquid phase separation of UBP1b of *Arabidopsis.* **a**, Protein structure (upper) and PrD-like score (lower) of UBP1b. **b**, *In vitro* phase separation assay. From top to bottom, GFP alone, GFP-tagged UBP1b full length protein and GFP-tagged UBP1b protein with prion-like domains deleted. The assay method was same as in Fig. 5. Scale bars, 10 μm. **c**, *In vitro* phase separation assay in the presence of *Arabidopsis* RNA. The assay was carried out using GFP-tagged UBP1b full length protein following the same method as in b. Total RNA extracted from Col-0 seedlings was supplemented for the amount indicated. Scale bars, 20 μm.

**Supplementary Fig. S5.**
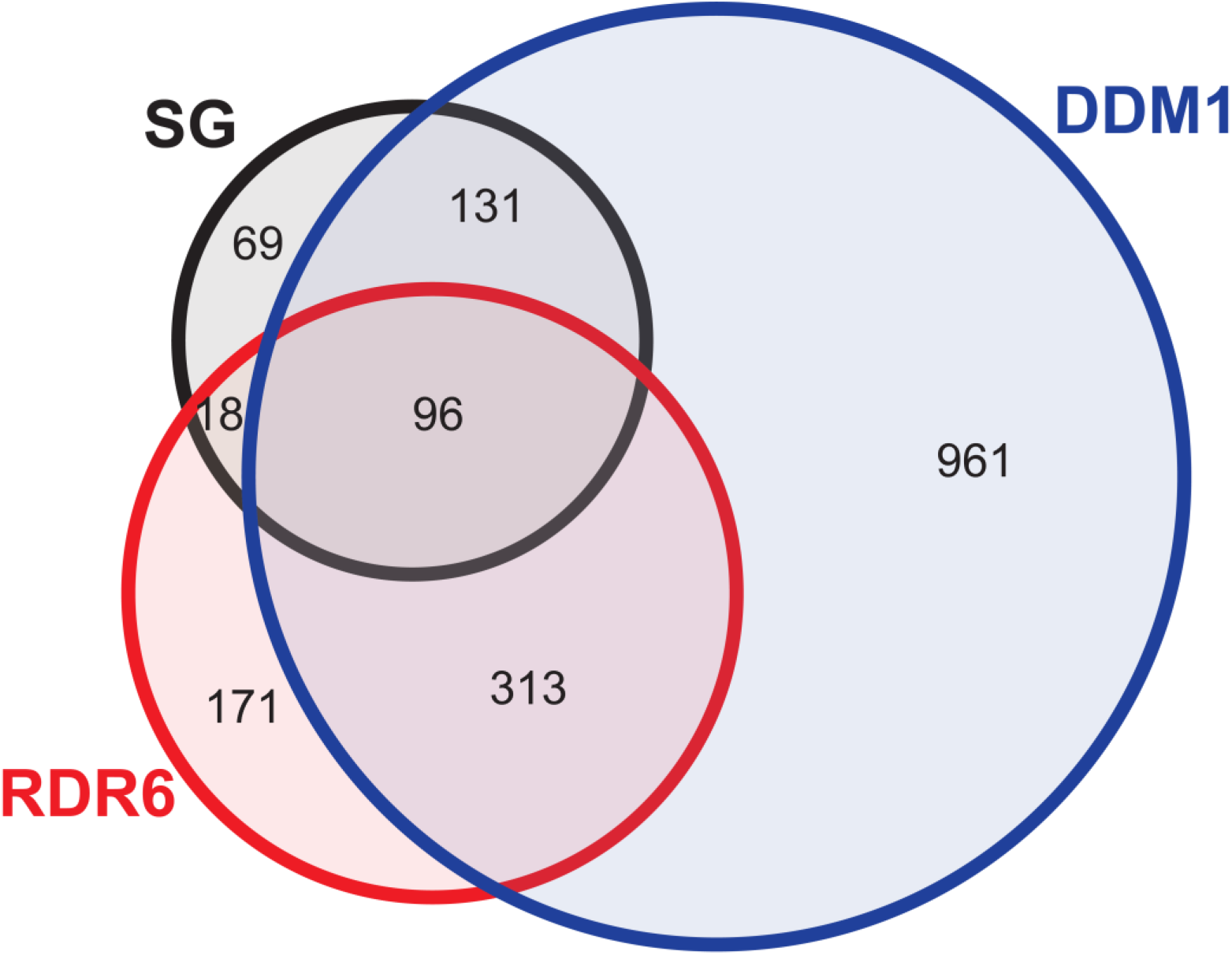
Overlap of SG-enriched transposons with those regulated by *RDR6* and *DDM1.* Venn diagram showing the transposons enriched in SG (black) and regulated by *RDR6* (red) and *DDM1* (blue) in *Arabidopsis.* The SG-enriched transposons are as determined in Fig. 6a. RDR6 targets are as identified in Fig. 3c. Transposons with increased expression in *ddm1* relative to the wildtype by at least two-fold were selected as DDM1 targets.

**Supplementary Fig. S6.**
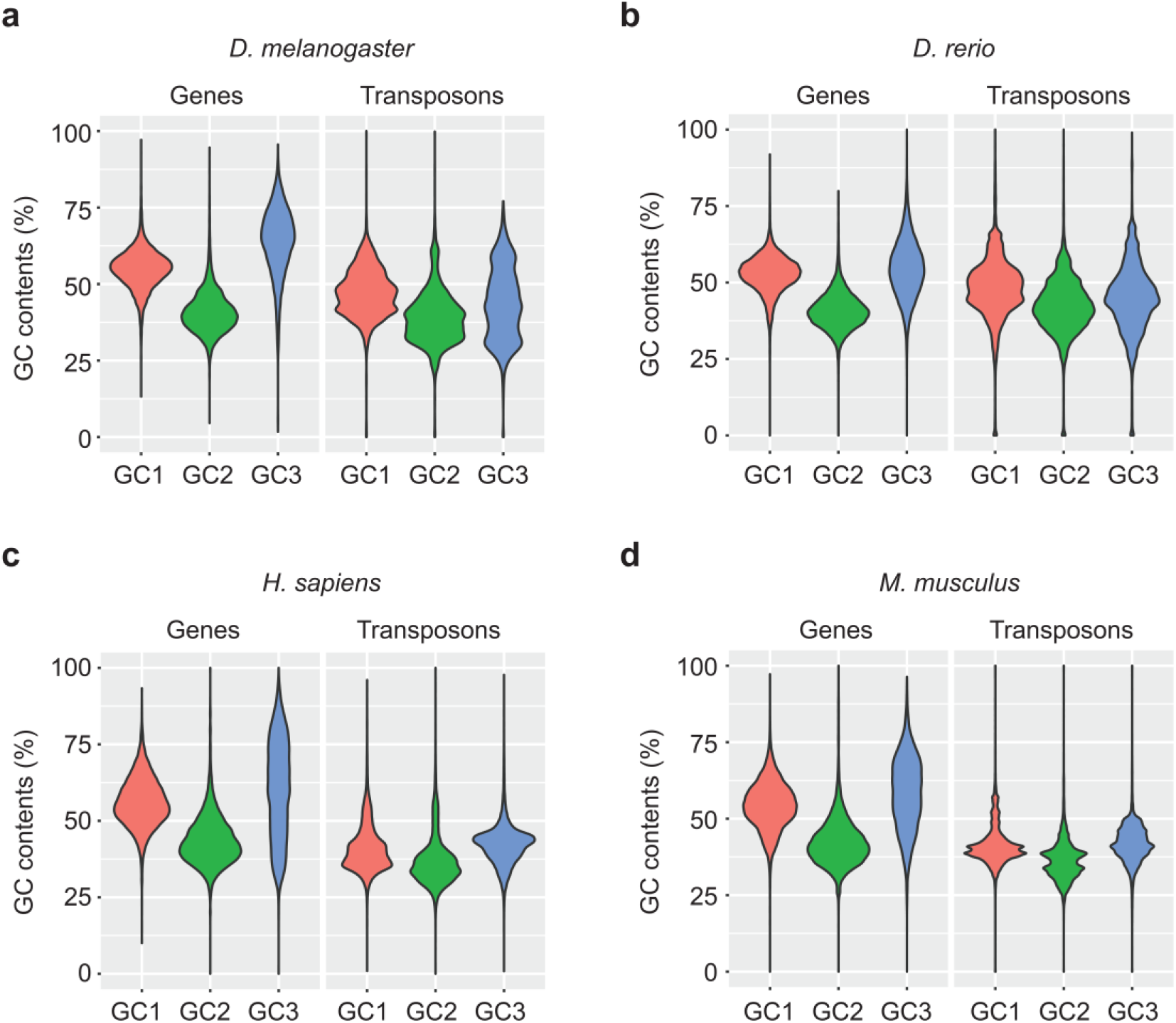
GC contents of genes and transposons in various species. GC1, GC2 and GC3 of fruit fly (**a**), zebrafish (**b**), human (**c**) and mouse (**d**) shown for genes and transposons separately.

**Supplementary Fig. S7.**
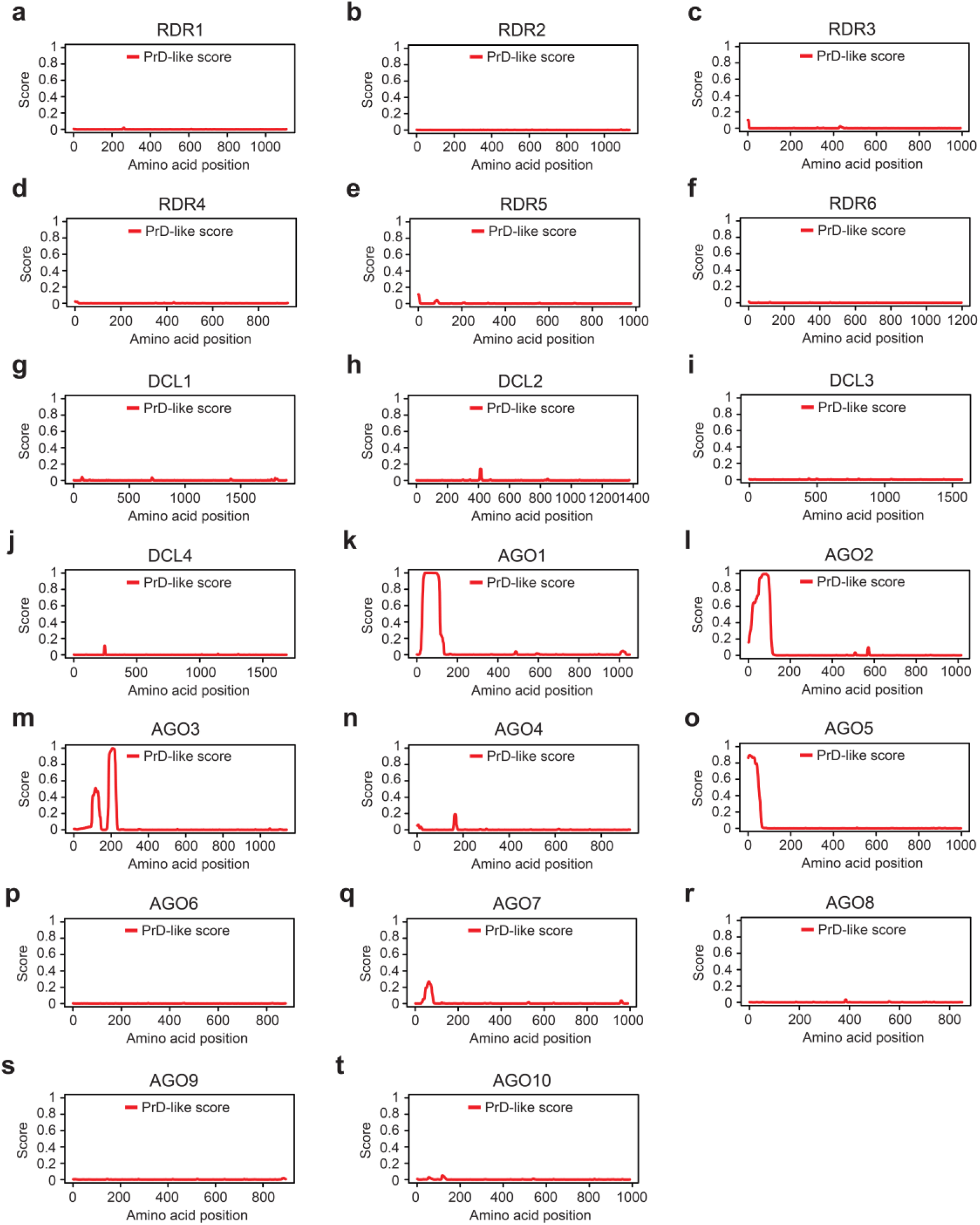
Prediction of prion-like domains in the siRNA biogenesis factors in *Arabidopsis.* Prion-like domains of RDR (**a-f**), DCL (**g-j**) and AGO (**k-t**) family proteins. The prediction was performed using the web-based tool PLAAC (http://plaac.wi.mit.edu/).

**Supplementary Table S1.**
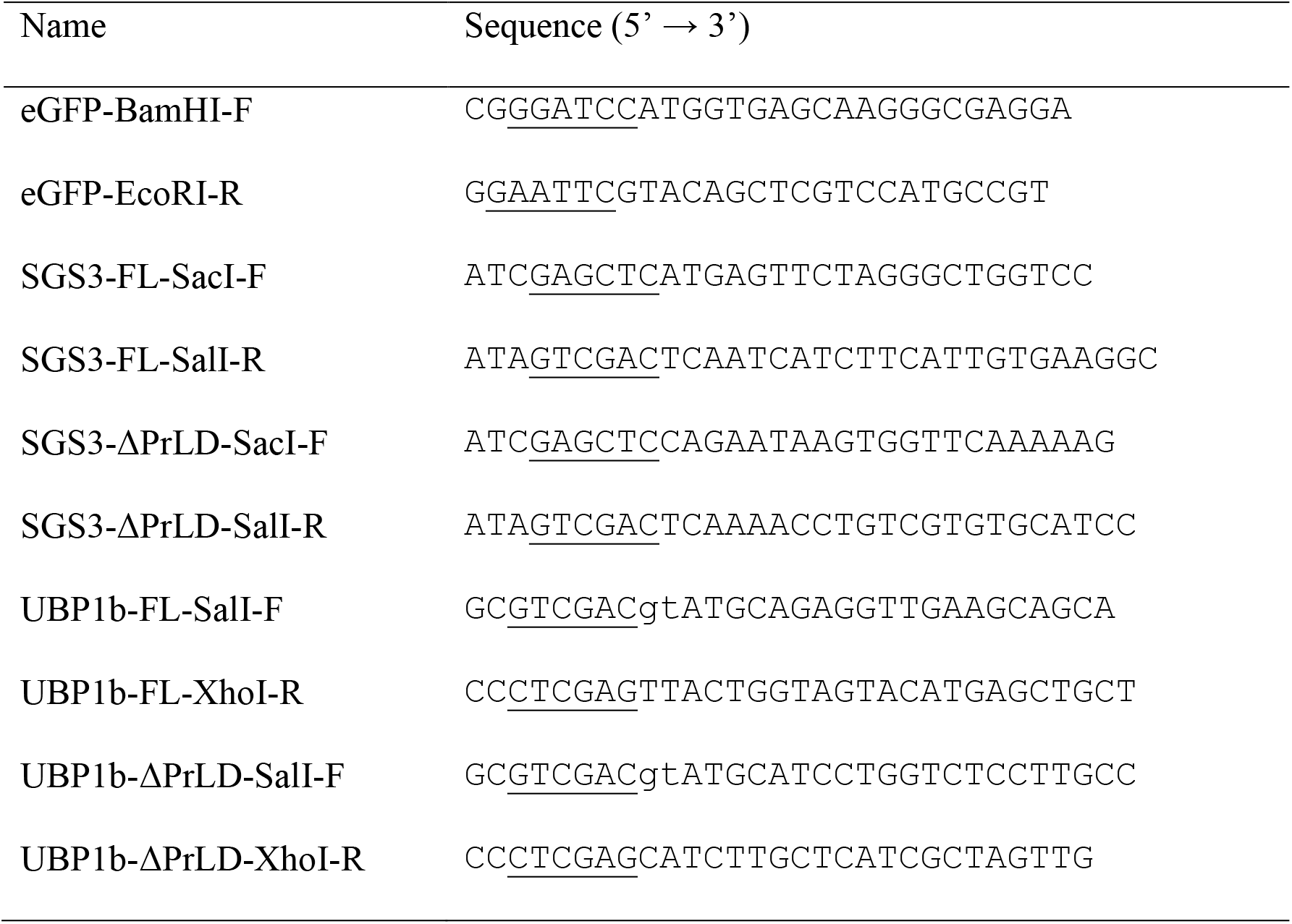
Oligonucleotides used in this study. The underlined is the restriction enzyme recognition sites.

**Supplementary Table S2.**
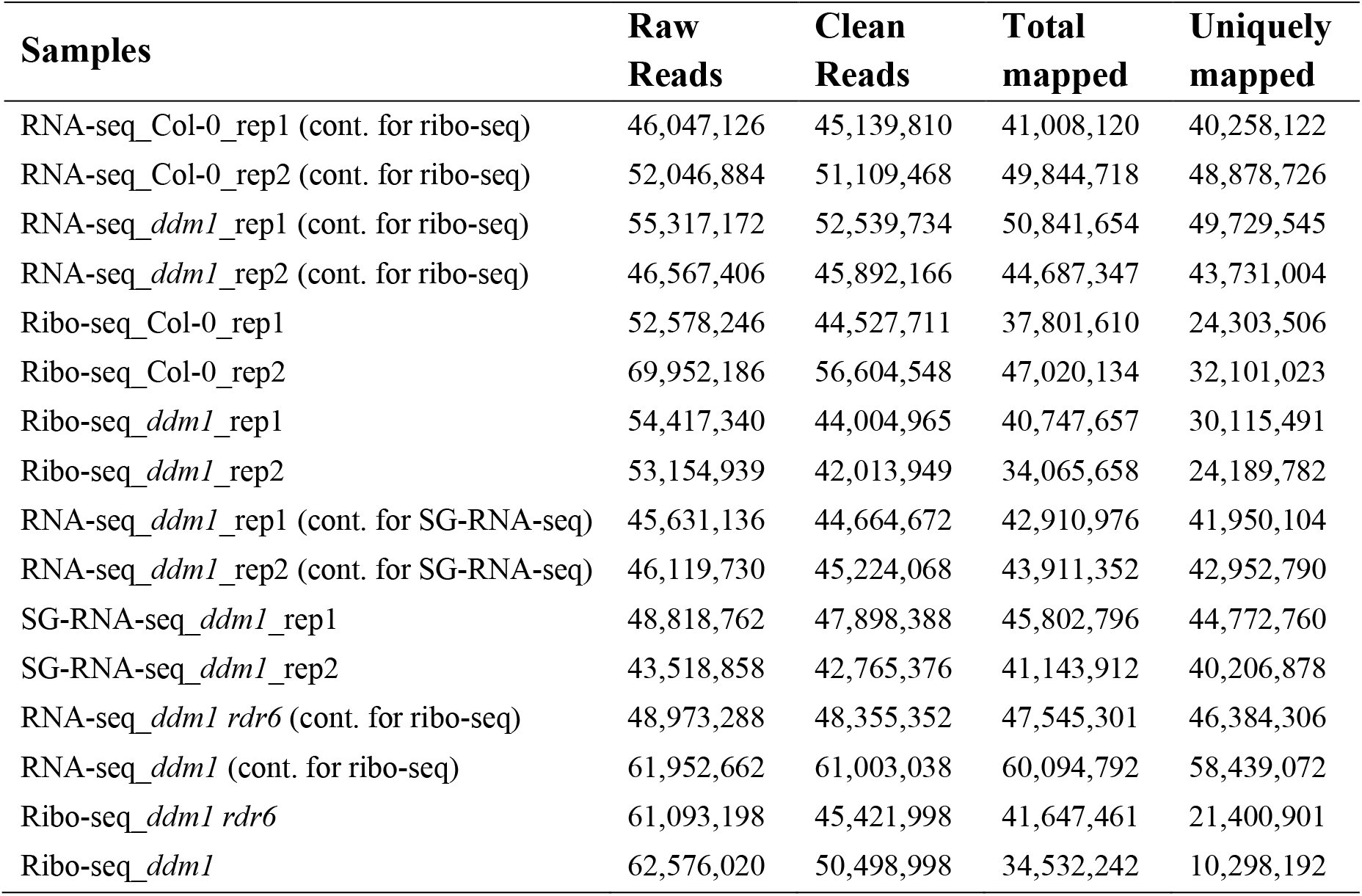
Summary of NGS data generated in this study. The NGS data generated in this study is publicly available under the accession code of PRJNA598331.

